# Fgf/Ets signalling in *Xenopus* ectoderm initiates neural induction and patterning in an autonomous and paracrine manners

**DOI:** 10.1101/2020.07.07.191288

**Authors:** Ikuko Hongo, Harumasa Okamoto

## Abstract

Fibroblast growth factor (Fgf) and anti-bone morphogenetic protein (Bmp) signals are derived from the organiser of mesoderm origin and cooperate to promote *Xenopus* neural development from the gastrula ectoderm. Using antisense oligos to Fgf2 and Fgf8 and dominant-negative Ets transcription factors, we showed that the expression of Fgf2, Fgf8, and Ets in ectoderm cells is essential to initiate neural induction both *in vivo* and *in vitro*. Our findings show that neural induction is initiated primarily by autonomous signalling in ectoderm cells, rather than by paracrine signalling from organiser cells. The signalling in ectoderm cells is transduced via the Fgf/Ras/Mapk/Ets pathway, independent of Bmp signal inhibition via the Fgf/Ras/Mapk/Smad1 route, as indicated by earlier studies. Through the same pathway, Fgfs activated position-specific neural genes dose-dependently along the anteroposterior axis in cultured ectoderm cells. The expression of these genes coincides with the establishment of the activated Ets gradient within the gastrula ectoderm. Organiser cells, being located posteriorly to the ectoderm, secrete Fgfs as gastrulation proceeds, which among several candidate molecules initially promote neural patterning of the induced neuroectoderm as morphogens.

**Summary statement:** Fgf/Ets signalling in ectodermal cells is required to initiate the expression of both anterior and posterior neural genes from the late blastula to gastrula stages, independent of anti-Bmp signalling.

## INTRODUCTION

Neural development in amphibians is initiated by the conversion of the intrinsic epidermal fate of the gastrula ectoderm to the anterior neural fate (i.e. neural induction). During subsequent development, the induced neuroectoderm of anterior identity is progressively patterned along the anteroposterior (AP) axis (i.e. neural patterning), yielding the fully differentiated central nervous system with the definitive AP pattern. Neural induction appears to be elicited by an instructive signal from the Spemann organiser towards the adjacent ectoderm during gastrulation; fibroblast growth factor (Fgf) and anti-bone morphogenetic protein (Bmp) signals are currently considered to mediate the action of the Spemann organiser (Stern, 2005). Loss-of-function and gain-of-function experiments have revealed that the combined activation of Fgf signalling and suppression of Bmp signalling in the ectoderm is necessary and sufficient to initiate neural induction (Wilson and Edlund, 2001; Stern, 2005). Bmps can promote the epidermal differentiation of ectoderm cells in an autocrine manner. The organiser cells inhibit this Bmp activity by releasing Bmp antagonists, such as Noggin and Chordin, which sequester Bmps from their receptors via direct binding, thereby suppressing the autonomous epidermal differentiation of ectoderm cells (Muñoz-Sanjuán and Brivanlou, 2002). Although organiser cells express Fgfs (Lea et al., 2009), their role in neural induction is undefined, since some previous experiments suggested that Fgfs derived from ectoderm cells primarily contribute to neural induction in an autonomous manner (Gruntz and Tacke, 1989; Sato and Sargent, 1989; Hongo et al., 1999; Delaune et al., 2005; Kuroda et al., 2005).

It is argued that active Fgf signalling reinforces the inhibition of Bmp signalling via the mitogen-activated protein kinase (Mapk)/Smad1 pathway (Pera et al., 2003; Kuroda et al., 2005). Mapk, a downstream transducer of the Fgf/Ras pathway, suppresses the transcriptional activity of Smad1, a downstream target of Bmp signalling, via the phosphorylation of inhibitory sites in the Smad1 linker region, leading to the cytoplasmic retention of Smad1 (Kretzschmar et al., 1997; Pera et al., 2003). This suggests that Fgf and anti-Bmp signals are integrated at the Smad1 level, which leads to adequate suppression of Bmp signalling in ectoderm cells to manifest their neural fate, which is the default state. However, some studies have suggested that Fgf signalling also functions in a Bmp-independent manner, exerting additional effects on ectoderm cells via pathways other than the Fgf/Ras/Mapk/Smad1 (Delaune et al., 2005; Marchal et al., 2009), but the mechanism remains to be clarified. One of the most promising candidates for such a pathway is the Fgf/Ras/Mapk/Ets pathway, whereby Fgfs can promote target gene expression by phosphorylating Ets family transcription factors via Mapk. We have previously shown that the Fgf/Ras/Mapk/Ets and anti-Bmp signalling pathways are integrated at the level of transcriptional regulation of *cdx4* (*Xcad3*), a posterior neural marker gene that is expressed at early-phase neural patterning (Haremaki et al., 2003). Activated Ets and Sox2, de-repressed via the inhibition of Bmp/Smad1 signalling (Mizuseki et al., 1998), cooperatively bind to their respective recognition sequences, activating *cdx4*. It is also notable that ascidians, which share the last common ancestor with vertebrates in chordate evolution, initiate neural induction via the Fgf/Ras/Mapk/Ets pathway (Bertrand et al., 2003; Miya and Nishida, 2003).

In the present study, we aimed to clarify the mode of action and the source of Fgf signals that are responsible for initiating neural induction and patterning. For this purpose, we blocked endogenous Ets activity in ectoderm cells using dominant-negative constructs of Ets, such as ΔxEts1 and ΔhElk1, and depleted several Fgf members in ectoderm cells using translation-blocking morpholino-modified antisense oligonucleotides (MOs). When we regard the activation of *otx2* and *sox2* as proving neural induction definitively, our *in vivo* and *in vitro* results provide evidence that the Ets transcriptional factor contributes to initiating neural induction via the Mapk/Ets route directly at the level of gene regulation. This is independent of Bmp signal inhibition at the level of Smad1 via the Mapk/Smad1 route. Since our *in vitro* results indicate the autonomous activity of ectoderm cells for neural induction, and Fgf4 has been suggested to function in neural induction (Marchal et al., 2009), we aimed to identify the members among a family of Fgf proteins that are expressed in ectoderm cells, exerting effects on them during neural induction. It turned out that Fgf2 and Fgf8 were primarily required for neural induction instead of Fgf4. We further attempted to verify the autonomous activity of ectoderm cells in Fgf signalling using isolated culture of ectoderm cells.

Finally, we asked about the role of Fgfs expressed in the organiser (Lea et al., 2009). There is an expression gradient of activated Ets within the gastrula ectoderm, with increasing levels towards the organiser, which is located posteriorly (Schohl and Fagotto, 2002). Here, we showed that Fgf/Ets signalling could activate position-specific neural genes, such as *otx2*, *hes7.1*, *foxb1*, and *cdx4*, dose-dependently along the AP axis in cultured ectoderm cells. The expression of these genes in normal development coincides with the establishment of the activated Ets gradient. This indicates that Fgfs derived from organiser cells contribute in a paracrine manner to initiating neural patterning of the induced neuroectoderm along the AP axis.

## RESULTS

### Plasmid construction for the expression of dominant-negative Ets transcription factors

To investigate the functional role of Ets transcription factors in neural induction and patterning, we prepared plasmids encoding several dominant-negative forms of Ets, as described in the materials and methods (Fig. 1A; ΔxEts1, ΔxEtv1, and ΔhElk1). They lack the activation domain, comprising mainly the DNA-binding Ets domain, thereby suppressing endogenous Ets transcriptional activity by competing for the target genes (Wasylyk et al., 1994). Two of the dominant-negative constructs were fused with the Engrailed repressor domain (EnR) to confirm that their native forms function as transcriptional activators (Fig. 1A; ΔhElk1·EnR and EnR· ΔxEtv1) (Conlon et al., 1996). ΔΔxEtv1 and EnR are used as negative control constructs.

**Fig. 1.**
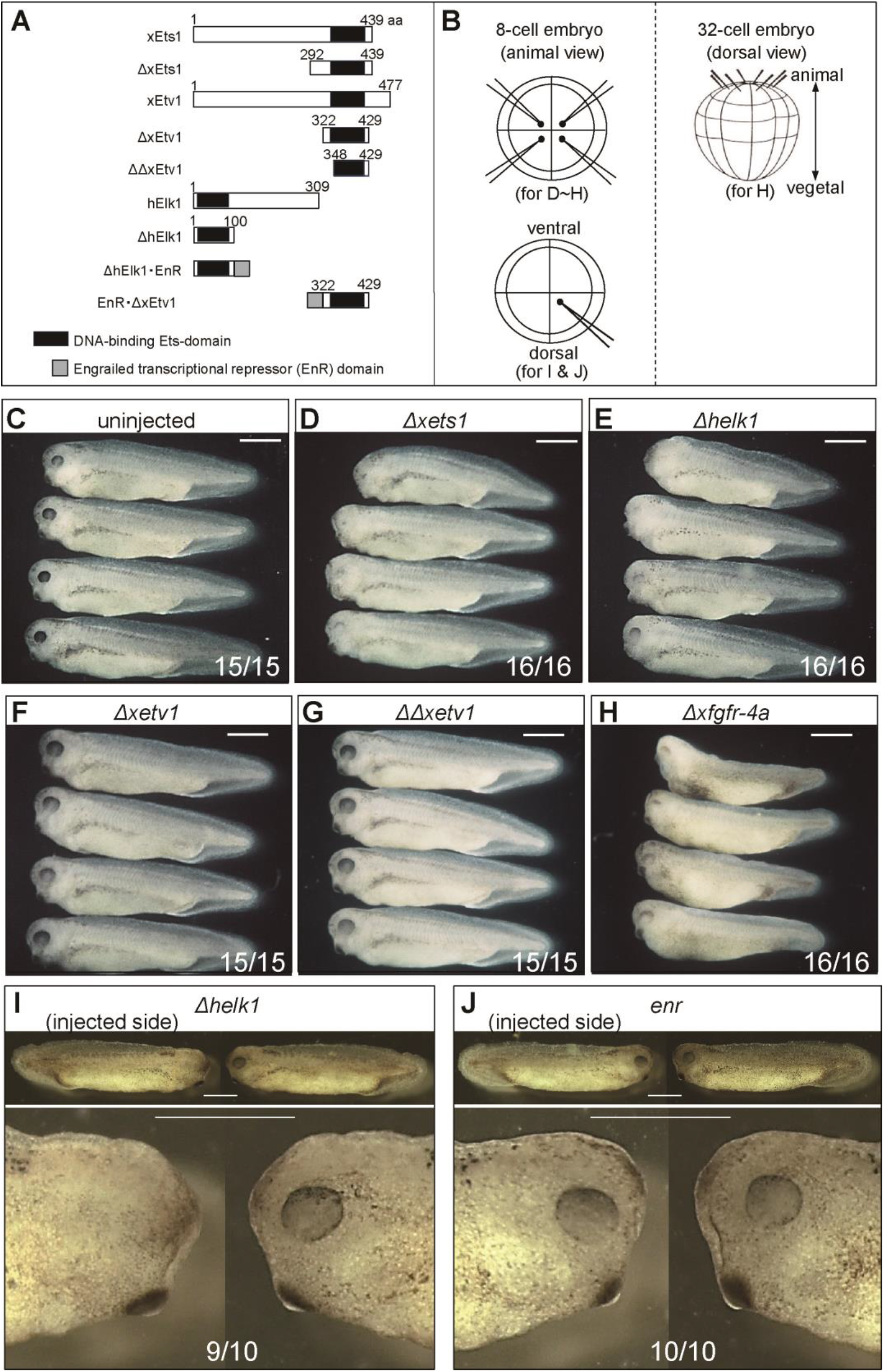
Suppression of anterior neural development *in vivo* by blocking Fgf/Ets signalling in ectoderm cells. (A) Structural features of Ets proteins and their dominant-negative derivatives. (B) Injection protocol. (C) Lateral views of uninjected embryos at stage 35/36. (D–J) Embryos were injected with synthetic RNA according to the respective protocol illustrated in (B), reared until stage 35/36, and photographed. Lateral views of embryos exhibiting typical phenotypes are shown. In (I) and (J), the lower panels are magnified versions of the upper ones, highlighting the anterior head region. Scale bar = 1 mm. The number of embryos exemplified in the photograph over the total number analysed is displayed on each panel.

### Ets transcriptional activity in ectoderm cells is required for anterior neural development

When synthetic *Δxets1* RNA (80 pg/blastomere) or *Δhelk1* RNA (40 pg/blastomere) was injected into four animal blastomeres at the 8-cell stage (Fig. 1B, upper left), morphological defects in the anterior region were prominently visible at the tadpole stage (Fig. 1D, E) compared with uninjected (Fig. 1C) or *ΔΔxetv1* RNA-injected (Fig. 1G; 80 pg/blastomere) controls. The defects included substantial reductions in head mass and eye tissue. Another dominant-negative construct, ΔxEtv1, was ineffective at the dose used (80 pg RNA/blastomere; Fig. 1F). However, ΔxEtv1, once fused with EnR, could effectively suppress Ets transcriptional activity (Fig. S1).

Upon injection of *Δhelk1* RNA (40 pg/blastomere) into a single dorsal-animal blastomere (Fig. 1B, lower left) to obtain a more targeted RNA distribution, the prominent defect was mostly confined to the eye tissue on the injected side (Fig. 1I). In embryos injected in either protocol, the axial structure appeared to have been minimally affected (Fig. 1D, E, I upper panels). This suggests that the anterior defects were elicited via direct effects on the ectoderm cells derived from RNA-injected blastomere(s) rather than secondarily, via neural-inducing organiser cells that generate primarily axial tissues. The tadpoles exhibiting severe anterior defects with a global RNA distribution of *Δxets1* or *Δhelk1* RNA (Fig. 1D, E) appeared to be phenotypic copies of tadpoles injected with *Δxfgfr-4a* RNA (Fig. 1H; Hongo et al., 1999; Hardcastle et al., 2000; Brunsdon and Isaacs, 2020) in a similar manner (Fig. 1B, right diagram). These experiments suggest that signalling through the Fgf/Ras/Mapk/Ets pathway in ectoderm cells is required for proper anterior neural development.

### Ets transcriptional activity in ectoderm cells is required to activate *sox2* and *otx2*

To ascertain whether Ets transcriptional activity was essential to initiate neural induction, we injected *Δhelk1* RNA with lineage tracer *gfp* RNA into three dorsal-animal blastomeres at the 16-cell stage (Fig. 2A). We examined the expression of the two early neural genes *sox2* (pan-neural marker) and *otx2* (anterior neural marker) using whole-mount *in situ* hybridisation. Both *sox2* and *otx2* are regarded as definitive markers for neural induction because their expression starts at the late blastula and early gastrula stages, respectively; the data for the temporal expression profiles of neural marker genes employed in this and the following experiments, except for *otx2* (Blitz and Cho, 1995), were based on previously published RNA-Seq analysis (Session et al., 2016). Typical results for *otx2* expression are shown in Fig. 2B and 2C. Dorsal view of embryos double-stained for *otx2* and *gfp* RNAs (Fig. 2C, left panels) showed a reduction in *otx2* expression in lineage-labelled dorsal ectoderm compared with uninjected controls (Fig. 2B, left panels). *Otx2* is expressed not only in a prospective anterior neural region of the outer ectoderm layer but also in the underlying mesoderm and endoderm layers during gastrulation (Blitz and Cho, 1995). To verify the site of *otx2* suppression, we bisected stained embryos sagittally through the dorsal–ventral mid-line (Fig. 2A, cut). Cut-surface view (Fig. 2C, middle and right panels) revealed that the tracer *gfp* RNA (stained red) was mostly confined to the ectoderm layer, and *otx2* expression was effectively suppressed in this tracer-labelled region. To ascertain the specificity of ΔhElk1, we co-injected wild-type *xets1* RNA to rescue *otx2* expression. Co-injection mostly recovered *otx2* expression, confirming that the suppression of *otx2* expression could be attributed to decreased Ets transcriptional activity in the ectoderm cells (Fig. 2D). We then examined the effects of ΔhElk1 on the expression of *sox2*, which encodes a transcriptional factor required to activate *otx2* (Mizuseki et al., 1998). The injection of *Δhelk1* RNA suppressed *sox2* expression, which was rescued by the co-injection of wild-type *xets1* RNA (Fig. 2E). In contrast, the expression of *chordin*, which marks the organiser tissue, was not affected by the injection of *Δhelk1* RNA (Fig. 2F). These results provide evidence that Ets transcriptional activity in ectoderm cells is essential for initiating neural induction.

**Fig. 2.**
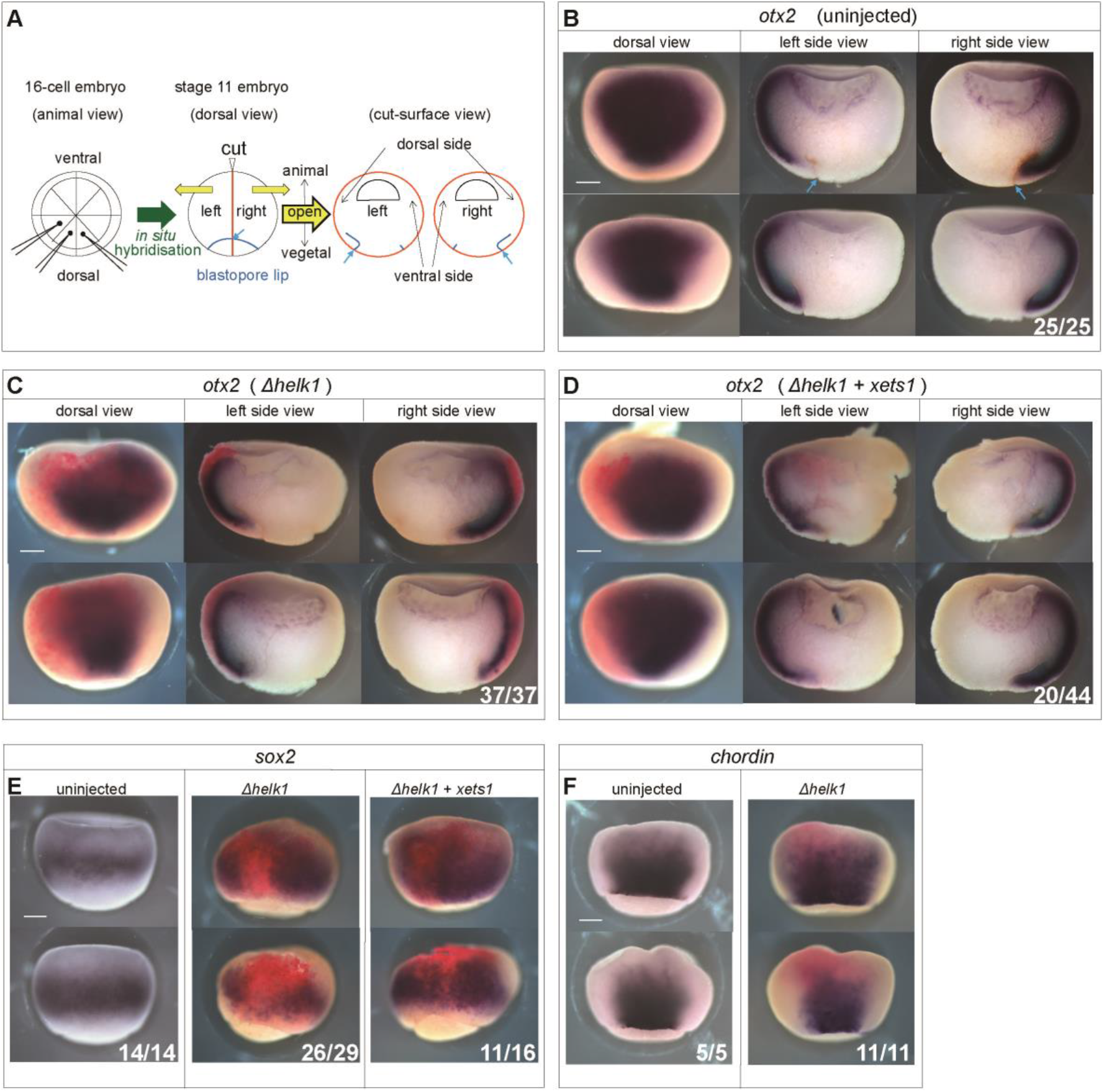
Suppression of *sox2* and *otx2* expression *in vivo* by blocking Ets transcriptional activity in ectoderm cells. (A) Experimental procedure highlighting the injection and bisection protocols. (B–F) Embryos were injected with *Δhelk1* RNA (45 pg/blastomere) plus the lineage tracer *gfp* RNA (20 pg/blastomere). Controls were not injected. In (D) and (E; right panel), wild-type *xets1* RNA (5.5 pg/blastomere) was co-injected. RNA *in situ* hybridisation was carried out on stage 11 gastrula embryos. Scale bar = 0.25 mm. The number of embryos exemplified in the photograph over the total number analysed is displayed on each panel.

### Ectoderm cells require Ets transcriptional activity to respond to Fgf signals

To confirm that ectoderm cells require Ets transcriptional activity for directly responding to Fgf signals, we exploited their primary culture in a microculture system as described by Kengaku and Okamoto (1995) (Fig. 3A). In the present study, the expression of a set of six position-specific neural marker genes was analysed using quantitative RT-PCR (Fig. 3), which differed from a set of genes used in the previous study. Ectoderm cells of the early gastrula were prepared from a combination of *Δxets1* RNA-injected embryos and *ΔΔxetv1* RNA-injected control embryos or a combination of *Δxfgfr-4a* RNA-injected embryos and uninjected control embryos. In the control experiments (Fig. 3B, D), Fgf induced ectoderm cells to express the six marker genes along the AP axis in a dose-dependent manner, similar to the previously described set of neural marker genes. Quantified data showed that lower doses of Fgf elicited more anterior marker genes, such as *bf1* (telencephalon marker), *rax* (eye marker), or *pax6* (telencephalon to spinal cord marker). In comparison, higher doses elicited more posterior marker genes, such as *en2* (midbrain-hindbrain boundary marker), *hoxc6* (whole spinal cord marker), or *cdx4* (posterior spinal cord marker) (Fig. 3F). We noted that the anterior marker genes were expressed in the absence of Fgf, but this default type of expression might be caused by the dissociation procedure during the preparation of ectoderm cells for culture (Gruntz and Tacke, 1989), which was necessarily employed in our culture system (Fig. 3A). The expression of the anterior neural genes peaked at a lower dose range of around 0.1 ng/mL and then declined as the dose increased (Fig. 3F and Fig. S2E, F, G). This characteristic of anterior neural genes being suppressed by a higher dose of Fgfs may explain why a high level of Fgf signalling notably suppressed anterior development, as previously reported (Cox and Hemmati-Brivanlou, 1995; Lamb and Harland, 1995; Pownall et al., 1996; Polevoy et al., 2019).

**Fig. 3.**
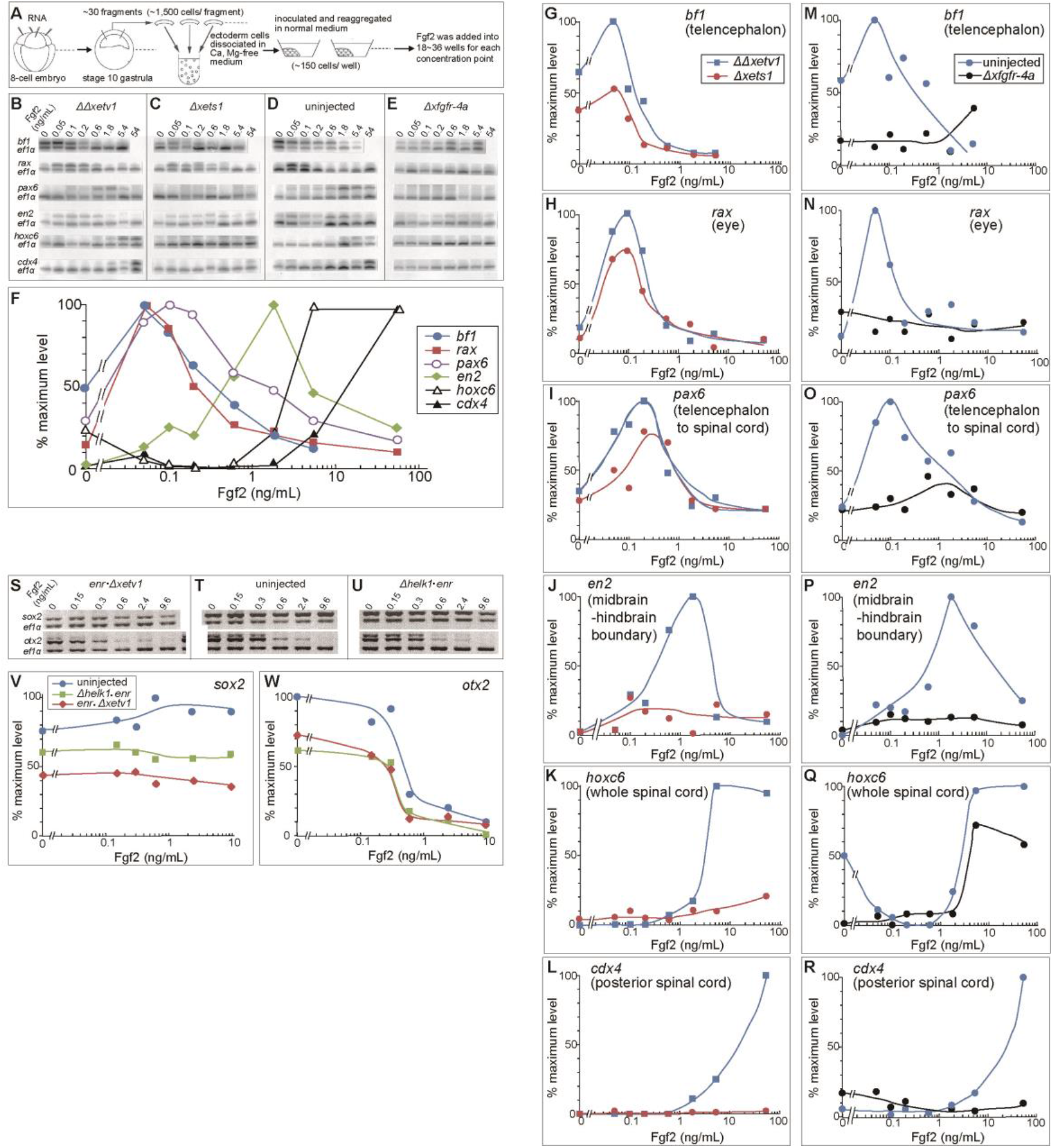
Suppression of Fgf-induced activation of position-specific neural marker genes in cultured ectoderm cells by blocking Ets transcriptional activity. (A) Experimental design for the primary culture of ectoderm cells. Cultured cells were harvested for mRNA isolation when control embryos reached stage 25 (for B–E) or stage 14 (for S–U). (B–E) PCR products of six position-specific neural marker gene mRNAs are indicated in (B), each co-amplified with *ef1α* mRNA (internal control). Ectoderm cells were prepared from embryos that had not been injected (D) or injected with *ΔΔxetv1* RNA (B), *Δxets1* RNA (C), or *Δxfgfr-4a* RNA (E) (63 pg/dorsal-animal blastomere and 105 pg/ventral-animal blastomere for each injection). (F) A quantitative comparison of the dose-response profiles of marker transcript levels. Data from control experiments (B) and (D) are combined. The ratio of the band intensities of the markers that of *ef1α* plotted against Fgf2 doses on a semi-log graph. The percentage of the maximum value of the ratio is presented in each profile. (G–L) Data from (B) and (C) are quantified and presented as in (F). (M–R) Data from (D) and (E) are quantified and presented as in (F). (S–U) PCR products of *sox2* and *otx2* mRNA, each co-amplified with *ef1α* mRNA (internal control). Ectoderm cells were prepared from embryos that had been injected with *enr·Δxetv1*RNA (S; 40 pg/dorsal-animal blastomere and 50 pg/ventral-animal blastomere) or *Δhelk1·enr* RNA (U; 4 pg/dorsal-animal blastomere and 5 pg/ventral-animal blastomere). For (T), ectoderm cells were prepared from uninjected embryos.

The overexpression of ΔxEts1, as well as ΔxFgfr-4a, suppressed the expression of all the above-mentioned marker genes (Fig. 3C,E, G–R), indicating that ectoderm cells required Ets transcriptional activity to activate neural genes in response to Fgf signalling. Notably, the anterior neural genes were less efficiently suppressed by ΔxEts1 than by ΔxFgfr-4a. A possible explanation for the incomplete suppression by ΔxEts1 is that it did not interfere with Smad1 function, while ΔxFgfr-4a did, since Ets transcription factors are downstream of the Fgf/Ras/Mapk pathway (Wasylyk et al., 1998). In ΔxEts1-overexpressing ectoderm cells, Mapk could boost the inhibition of Bmp signalling via Smad1 phosphorylation at a level sufficient to activate anterior neural genes to a certain extent, possibly combined with residual weak Mapk/Ets signalling. In contrast, ΔxEts1 efficiently suppressed the expression of posterior neural genes elicited by the increased dose of Fgf, suggesting that the reinforcement of Bmp signal inhibition via the Mapk/Smad1 route was not necessarily essential for posterior neural gene expression as long as Fgf signalling levels were kept high.

### Fgf/Ets signalling activates *sox2* and *otx2* in ectoderm cells

To establish the direct role of the Ets transcription factor in ectoderm cells in initiating neural induction in response to Fgf signals, we analysed the expression of *sox2* and *otx2* in cultured ectoderm cells with or without dominant-negative Ets expression. In this series of experiments, we employed ΔhElk1·EnR and EnR·ΔxEtv1 (Fig. 1A) to ascertain their native functions as transcriptional activators. EnR·ΔxEtv1 overexpression in ectoderm cells could effectively suppress *pax6* expression (Fig. S1), even more strongly than ΔxEts1 overexpression could (Fig. 3I). In contrast, ΔhElk1·EnR had little effect on *pax6* expression. The difference in the suppression efficiency among these three dominant-negative constructs may reflect the difference in DNA-binding specificity of their Ets domain.

The overexpression of ΔhElk1·EnR or EnR·ΔxEtv1 could suppress the expression of both *sox2* and *otx2* (Fig. 3S–W), although the degree of suppression was not prominent, being similar to the degree of suppression of other anterior neural marker genes, which were expressed later in neural development, by ΔxEts1 (Fig. 3G–I). The incomplete suppression of *sox2* and *otx2* might be explained in the same way as that of the other anterior neural marker genes by ΔxEts1, as noted before. These results imply that Fgf signalling in ectoderm cells contributed to neural induction at the level of transcriptional activation of target genes via the Mapk/Ets route, independent of Bmp signalling inhibition via the Mapk/Smad1 route.

### Fgf2 and Fgf8 expression in ectoderm cells promotes anterior neural development via the Fgf/Ets pathway

To search for Fgf ligands that promote neural induction and their source, we investigated the phenotypic effects of reducing Fgf2, Fgf3, Fgf4, Fgf8, and Fgf9 expression in ectoderm cells using respective translation-blocking antisense MOs. Transcripts encoding these Fgfs are present in embryos during the late blastula and early gastrula stages based on previous RNA-Seq data (Session et al., 2016). Among them, Fgf4 alone was examined and indicated to contribute to neural induction (Marchal et al., 2009). Upon injection of each Fgf MO into a single dorsal-animal blastomere at the 8-cell stage, however, we found that Fgf2 MO and Fgf8 MO were much more potent than the other MOs, including Fgf4 MO, in causing anterior head defects, such as reductions in head mass and eye tissue on the injected side (Fig. 4A, C, E–G; H for collected data). The morphological phenotypes elicited by Fgf2 MO and Fgf8 MO were quite similar to those induced by overexpressed dominant-negative Ets (Fig. 1D, E, I) or xFgfr-4a (Fig. 1H) in ectoderm cells in that they exhibited prominent anterior head defects, ensuring that the overall axial structure remained well preserved. These apparent phenotypic similarities imply that the Fgf/Ets pathway functions autonomously in ectoderm cells for anterior neural development, given that Fgf MOs were targeted primarily to ectoderm cells. We obtained several lines of evidence supporting the specificity of Fgf2 MO and Fgf8 MO. Most embryos injected with 5mis-Fgf2 MO or 5mis-Fgf8 MO formed a normal anterior head structure (Fig. 4B, D, H). The inhibition of anterior head development in each Fgf2 and Fgf8 morphant was primarily rescued by the co-injection of either cognate *fgf2* or *fgf8* RNAs (Fig. 4I–N). Most significantly, co-injection of wild-type *Xenopus etv1* (*xetv1*) RNA could also recover anterior head development (Fig. 4O, P), ruling out the possibility of non-specific side effects of the MOs, such as deteriorative effects on embryonic morphogenesis via an innate immune response (Gentsch et al., 2018; Paraiso et al., 2019).

**Fig. 4.**
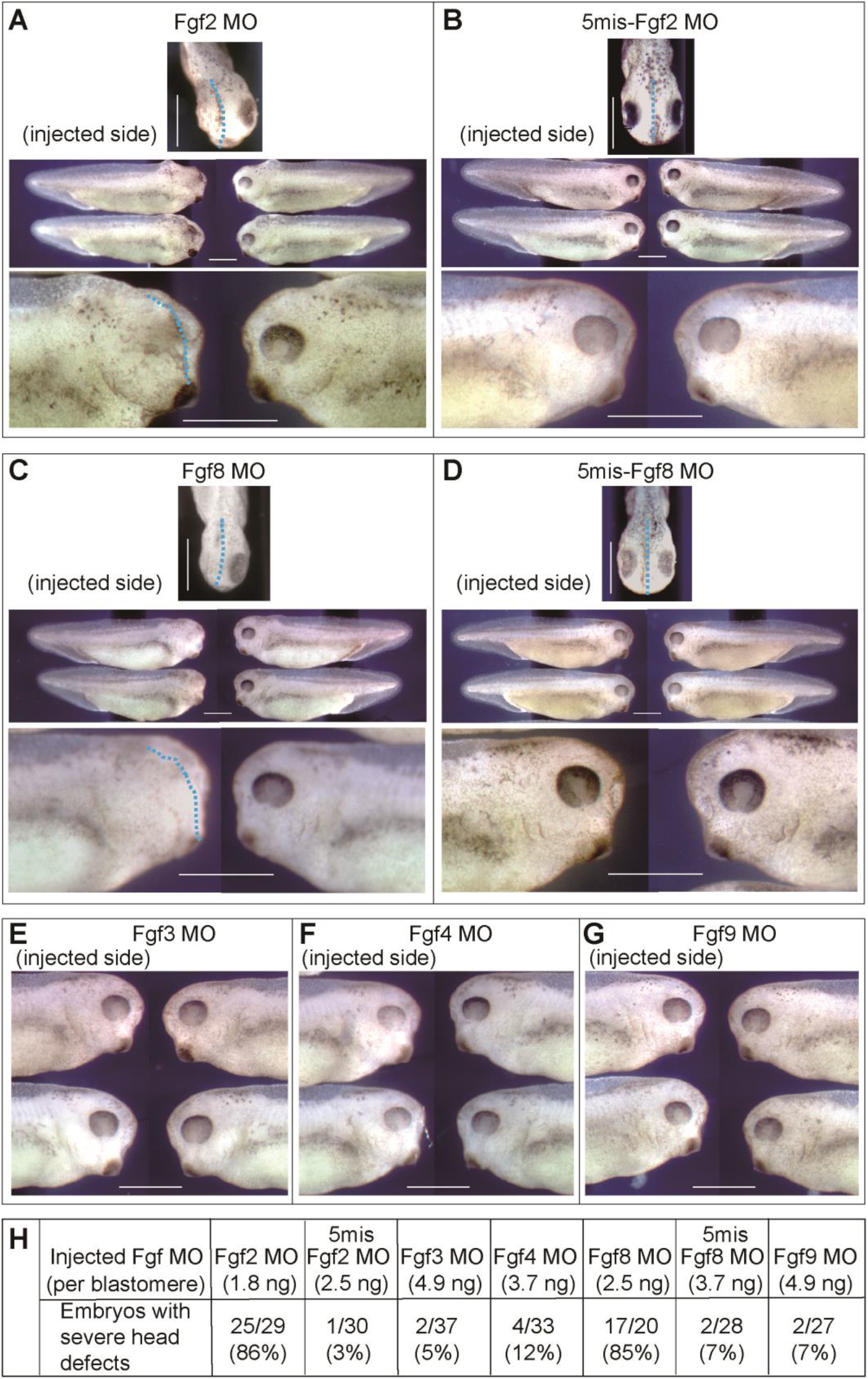
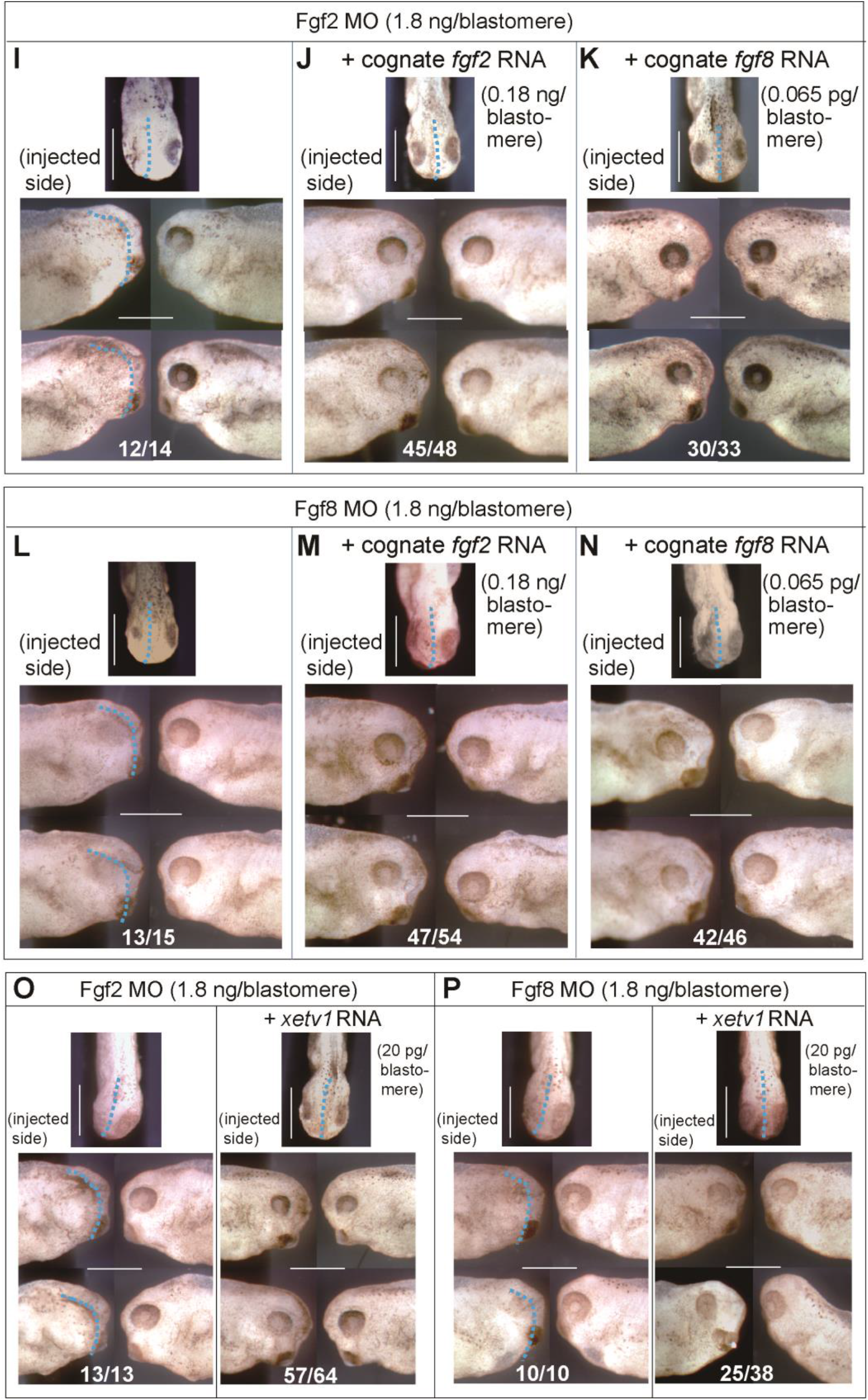
Suppression of anterior head development *in vivo* by depleting Fgf2 or Fgf8 in ectoderm cells. Antisense morpholino oligo (MO) or 5mis-MO was injected into a single dorsal-animal blastomere at the 8-cell stage. Injected embryos were cultured until stage 35/36. Scale bar = 1 mm. (A–D) Upper panels show a dorsal view of the anterior head of an injected embryo. Middle panels show a lateral view of the whole body of two injected embryos. Lower panels show a lateral view of the anterior head region of another injected embryo at a higher magnification. Broken lines in the upper and lower panels represent the assumed mid-line of affected embryos. (E–G) Lateral views of the anterior head region of two injected embryos are shown. (H) Numerical data were collected from the experiments exemplified in (A–G) and displayed for comparison. (I) Fgf2 MO was injected with no additives, (J) with cognate *fgf2* RNA, or (K) cognate *fgf8* RNA. (L) Fgf8 MO was injected with no additives, (M) with cognate *fgf2* RNA, or (N) cognate *fgf8* RNA. (O) Fgf2 MO or (P) Fgf8 MO was injected without (left panels) or with (right panels) wild-type *xetv1* RNA. In (I–P), the number of embryos exemplified in the photograph over the total number analysed is displayed on each panel.

### Autonomous Fgf/Ets signalling in ectoderm cells is required for neural induction

Depleting Fgf2 or Fgf8 in ectoderm cells by injecting the respective antisense MO into a single dorsal-animal blastomere at the 8-cell stage (Fig. 5A) reduced the expression of *otx2* and *sox2* in the prospective neural region of the stage 11 gastrula on the injected side compared with 5mis-Fgf2 MO controls (Fig. 5B, C). In contrast, the expression of *chordin* in the organiser region was not affected (Fig. 5D). These results provide evidence that Fgf2 and Fgf8 in ectoderm cells function autonomously during neural induction. The suppression of *otx2* and *sox2* expression in Fgf morphants could be efficiently rescued by the co-injection of the cognate *fgf* RNA (Fig. 5E, F left panels) or wild-type *xetv1* RNA (Fig. 5E, F, right panels). The rescue by a wild-type Ets protein strongly supports our claim that neural-inducing Fgfs act via the Fgf/Ras/Mapk/Ets pathway.

**Fig. 5.**
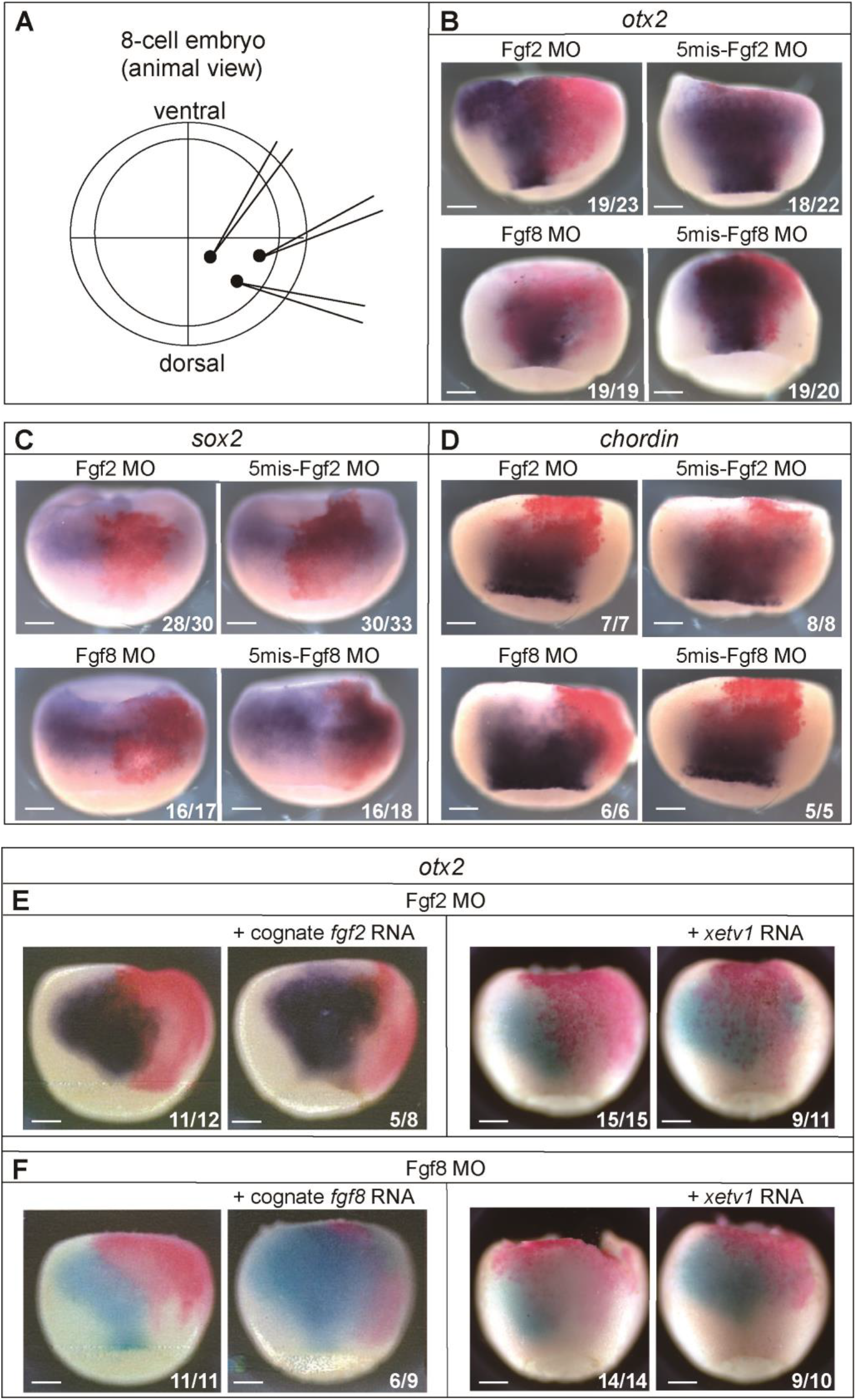
Suppression of *otx2* and *sox2* expression *in vivo* by depleting Fgf2 or Fgf8 in ectoderm cells. (A) Injection protocol. We poked a single dorsal-animal blastomere at three separate sites to obtain a more global MO distribution in the progeny of the injected blastomere. (B–D) The upper panel embryos were injected with Fgf2 MO (left panel) or 5mis-Fgf2 MO (right panel) at a dose of 1.8 ng/blastomere, while embryos in the lower panels were injected with Fgf8 MO (left panel) or 5mis-Fgf8 MO (right panel) at the same dose. (E) Embryos in the left panels were injected with Fgf2 MO (2.4 ng/blastomere) with or without cognate *fgf2* RNA (0.15 ng/blastomere), while embryos in the right panels were injected with Fgf2 MO (2.4 ng/blastomere) with or without wild-type *xetv1* RNA (20 pg/blastomere). (F) In the left panels, Fgf8 MO (2.4 ng/blastomere) was injected with or without cognate *fgf8* RNA (0.065 pg/blastomere), while in the right panels, Fgf8 MO (2.4 ng/blastomere) was injected with or without wild-type *xetv1* RNA (20 pg/blastomere). Scale bar = 0.25 mm. The number of embryos exemplified in the photograph over the total number analysed is displayed on each panel.

We further verified the autonomous activity of ectoderm cells in Fgf signalling by culturing them in isolation. Under Fgf-free culture conditions, we found that ectoderm cells expressed *fgf8* in addition to anterior neural genes. The time course of *fgf8* activation was roughly parallel with that of anterior neural genes, such as *pax6* and *rax*, whereas *otx2* expression decreased gradually to a certain level during this period (Fig. 6A). As shown before (Fig. 3W, uninjected control), *otx2* expression in cultured ectoderm cells was gradually suppressed as the added Fgf ligand dose increased. Thus, the temporal expression patterns shown in Fig. 6A could be explained by ectoderm cells synthesising and secreting Fgf8 gradually to either activate or suppress anterior neural genes. When ectoderm cells were prepared from Fgf8 morphants, the expression of *fgf8* and other neural genes, including *sox2*, was reduced compared to ectoderm cells prepared from control embryos injected with 5mis-Fgf8 MO (Fig. 6B). The expression of the same set of genes could also be suppressed by pharmacological inhibitors of the Fgf/Ras/Mapk pathway, such as SU5402 (against Fgf receptor) and U0126 (against Mapk kinase), albeit to a varying degree (Fig. 6C). Finally, the overexpression of EnR·ΔxEtv1 or ΔhElk·EnR in ectoderm cells also reduced the expression of these genes (Fig. 6D). Collectively, autocrine Fgf8 signalling in ectoderm cells contributed to neural induction via the Fgf/Ras/Mapk/Ets pathway.

**Fig. 6.**
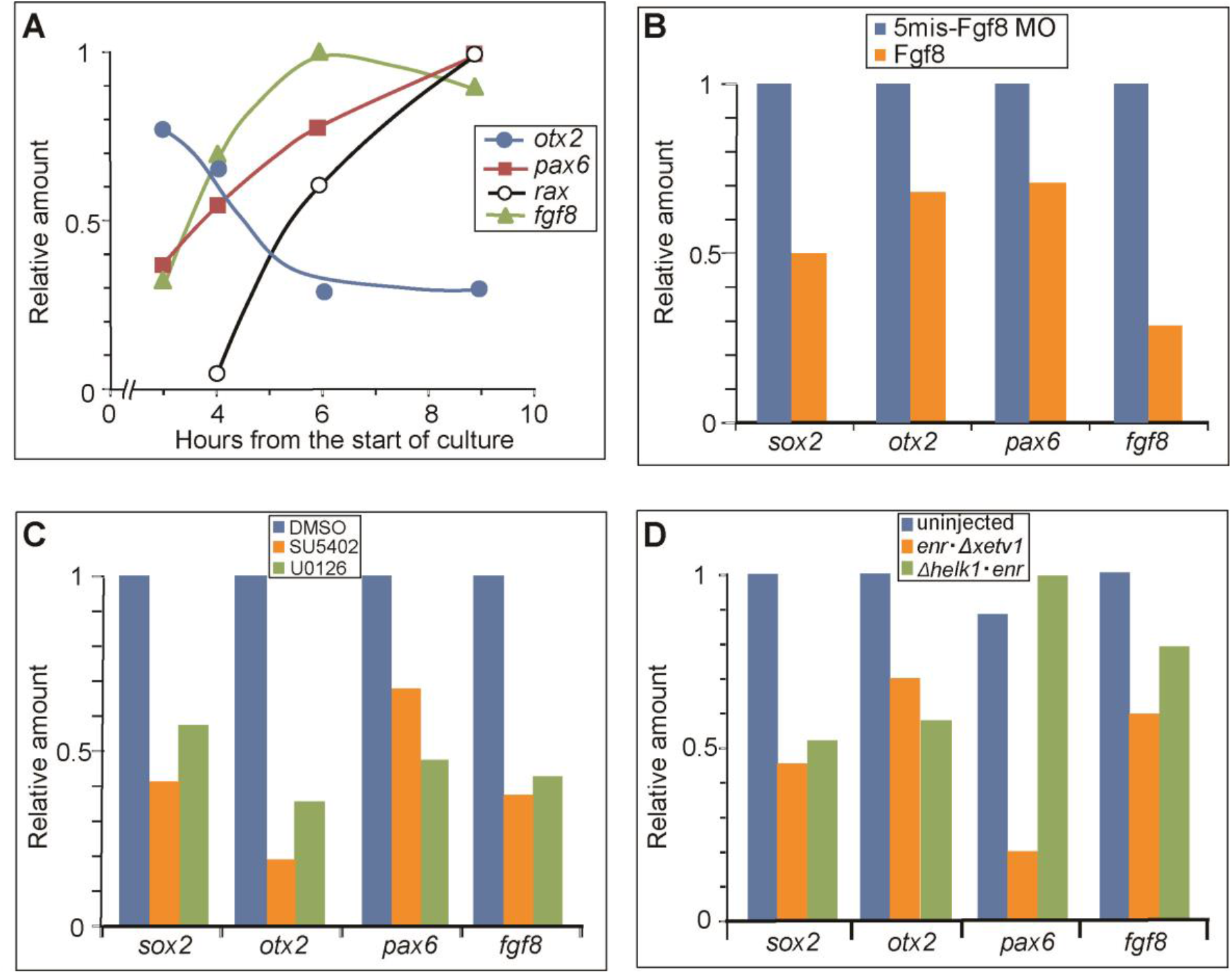
Fgf8/Ras/Mapk/Ets signalling in cultured ectoderm cells activates *fgf8*, *sox2,* and anterior neural genes in an autocrine manner. (A) Time course of *fgf8* and anterior neural gene activation in cultured ectoderm cells. Ectoderm cells were prepared from stage 10 gastrula embryos and cultured without Fgf. They were harvested serially after the start of the culture and processed for RT-PCR. Relative amounts of transcripts from each gene are plotted against the time of harvest. (B–D) Suppression of autonomous *fgf8* and neural gene activation in ectoderm cells by depleting Fgf8 (B) by adding inhibitors of the Fgf/Ras/Mapk pathway (C) or by blocking Ets transcriptional activity (D). (B) Ectoderm cells were prepared from stage 10 embryos that had been injected with Fgf8 MO or 5mis-Fgf8 MO (4.8 ng/dorsal-animal blastomere and 6.0 ng/ventral-animal blastomere) at the 8-cell stage, as described in Fig. 3A. (C) Ectoderm cells were prepared from intact stage 10 embryos and cultured in the presence of SU5402 (400 μM) or U0126 (200 μM). (D) Ectoderm cells were prepared from stage 10 embryos that had been injected with *enr·Δxetv1* RNA (40 pg/dorsal-animal blastomere and 50 pg/ventral-animal blastomere) or *Δhelk1·enr* RNA (4 pg/dorsal-animal blastomere and 5 pg/ventral-animal blastomere) at the 8-cell stage. After culturing without the addition of Fgf until control sib-embryos reached stage 14, they were harvested and subjected to RT-PCR. The band intensity was analysed using ImageJ. Relative amounts of transcripts from each gene were compared.

### Fgf/Ets signalling in ectoderm cells promotes neural patterning as an early morphogenic factor

To verify the role of Fgf/Ets signalling in initiating neural patterning, we examined the expression profiles of position-specific neural marker genes of the early developmental stage in ectoderm cells cultured with Fgf. They include *otx2* (forebrain marker), *hes7.1* (midbrain– hindbrain boundary marker), *foxb1* (hindbrain marker), and *cdx4* (posterior spinal cord marker), which are first activated around stage 10, peaking around stage 12 in normal development (Session et al., 2016). Their expression patterns in whole embryos at stage 11, as revealed by *in situ* hybridisation, are presented in Fig. 7A, with that of *sox2*. It should be noted that the expression profiles of position-specific marker genes used earlier (Fig. 3F), except *cdx4*, did not necessarily reflect the initial phase of neural patterning, since their expression began during the mid-gastrula stage (around stage 12) and reached a peak during the early neurula stage (around stage 15) in normal development.

**Fig. 7.**
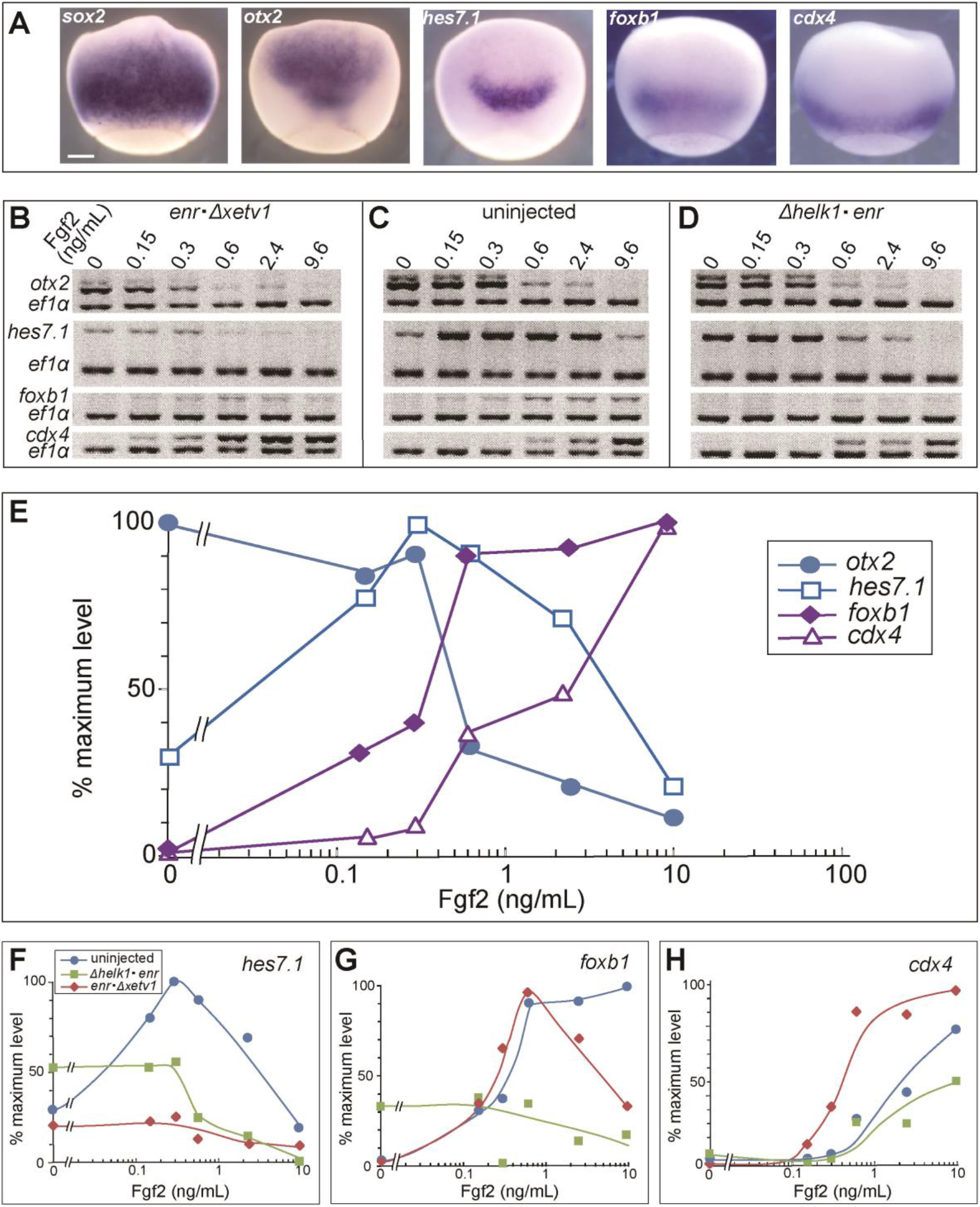
Fgf/Ets signalling in cultured ectoderm cells activates position-specific early neural marker genes in a dose-dependent manner. (A) Expression pattern of *sox2* and the position-specific early markers at stage 11. Scale bar = 0.25 mm. (B–D) The experimental design was as described in Fig. 3A. *enr·Δxetv1* RNA and *Δhelk1·enr* RNA were injected as in Fig 6D. Cultured cells were harvested when control embryos reached stage 14. The band intensities were analysed using ImageJ. (E) A quantitative comparison of the dose-response profiles of transcript levels of the four markers. Data in (C) were quantified. (F–H) Suppression of Fgf-induced activation of the position-specific early markers in cultured ectoderm cells by ΔhElk1·EnR or EnR·ΔxEtv1. Data in (B), (C), and (D) were quantified.

Control experiments showed that Fgfs could induce ectoderm cells to express the earlier marker genes along the AP axis in a dose-dependent manner (Fig. 7C, E; Fig. S2A, B, D), similar to the later phase marker genes shown in Fig. 3F. The overexpression of EnR·ΔxEtv1 or ΔhElk1·EnR reduced the expression of the marker genes to a varying degree. EnR·ΔxEtv1 effectively suppressed *hes7.1* expression but had little effect on *foxb1* and even enhanced *cdx4* expression, whereas ΔhElk1·EnR effectively suppressed both (Fig. 7F–H). *Otx2* expression was reduced by the two constructs to a similar extent (Fig. 3W). The varying degree of suppression may be interpreted as the difference in DNA-binding specificities of the respective Ets domains of xEtv1 and hElk1 to each marker gene, as noted before. Our data suggest that Fgf signalling in ectoderm cells promoted neural patterning as early as the early gastrula stage in a dose-dependent manner, which required Ets proteins as transcriptional activators.

## DISCUSSION

### Fgf2 and Fgf8/Ets signalling in ectoderm cells is required to initiate neural induction

By overexpressing a dominant-negative form of an Ets transcription factor (ΔhElk1), we showed that endogenous Ets transcriptional activity in ectoderm cells is specifically required to initiate neural induction, as revealed by the suppression of *sox2* and *otx2* expression and its rescue by wild-type xEts1 in gastrula embryos (Fig. 2C, D, E). We also showed the requirement of Ets transcriptional activity for cultured ectoderm cells to activate *sox2* and *otx2* (Fig. 3V, W). Our previous studies, in which ΔxFgfr-4a was overexpressed in cultured ectoderm cells, showed that Fgf receptor signalling is required for ectoderm cells to adopt a neural fate in response to co-cultured intact organiser cells (Hongo et al., 1999). Furthermore, the morphological phenotype of tadpoles elicited by ΔhElk1 or ΔxEts1 was quite similar to that by ΔxFgfr-4a (Fig. 1D, E, H). These findings indicate that the Fgf/Ras/Mapk/Ets pathway in ectoderm cells is essential for initiating neural induction.

Numerous *fgf* transcripts are present in blastula and gastrula embryos, as shown by RNA-Seq analysis (Suzuki et al., 2017). A report suggested that Fgf4 is a member of the Fgf protein family that functioned in neural induction (Marchal et al., 2009), but the possibility of contributions of other Fgf members remains unaddressed. We attempted to identify members of the Fgf family exerting effects on ectoderm cells during neural induction. For this, we knocked down Fgf activity in ectoderm cells by injecting several translation-blocking MOs into a single 8-cell-stage dorsal-animal blastomere. The depletion of Fgf2 and Fgf8 in ectoderm cells by antisense MO most effectively suppressed *sox2* and *otx2* expression in gastrula embryos (Fig. 5B, C) and caused severe defects in the anterior head in tadpoles (Fig. 4A, C), similar to the effects of the overexpression of dominant-negative Ets proteins (Fig. 1D, E; Fig. 2C, E middle panels). The suppression of *otx2*, as well as anterior head development, in Fgf morphants was rescued by xEtv1, a wild-type *Xenopus* Ets protein (Fig. 4O, P; Fig. 5E, F, right panels). Collectively, these results provide evidence that Fgf2 and Fgf8 act via the Fgf/Ras/Mapk/Ets pathway in ectoderm cells, independent of the Mapk/Smad1 branch, to initiate neural induction, considering the expression of *sox2* and *otx2* as an indication of the start of neural induction.

### Fgf2 and Fgf8/Ets signalling acts autonomously in neural induction

The microinjection of dominant-negative forms of Ets or Fgf MOs into animal blastomeres at the 8- or 16-cell stage exhibited minimal effects on the functional activities of the organiser cells, since the expression of *chordin* at the gastrula stage (Figs 2F, 5D), and subsequent formation of the axial structures were barely affected (Fig. 1D, E; Fig. 4A, C). This indicates that neural-inducing Fgfs function autonomously between ectoderm cells rather than in a paracrine manner from the organiser cells. Consistently, some previous experiments suggested the possible involvement of ectoderm cells in neural induction, which contribute in an autonomous manner (Delaune et al., 2005; Kuroda et al., 2005). The idea is based on the observation that ectoderm cells subjected to prolonged dissociation differentiate into neural cells of anterior identity in the absence of organiser cells (Gruntz and Tacke, 1989; Sato and Sargent, 1989) and that this autonomous type of *in vitro* neuralization is blocked by ΔxFgfr-4a overexpression (Hongo et al., 1999).

In the microculture system we used, a dissociation procedure during the preparation of the ectoderm cells was a prerequisite step, which caused the expression of pan-neural *sox2* and anterior neural marker genes in the absence of Fgf (Fig. 3G, H, I, M, N, O, V, and W). We found in this culture system of ectoderm cells that *fgf8* was also activated autonomously along with the neural genes (Fig. 6A). In normal development, *fgf8* is activated as early as the late blastula (stage 9), and maternally expressed *fgf2* is also present during the blastula and gastrula stages (Suzuki et al., 2017). Our preliminary experiments showed that Fgf2 MO could suppress the expression of *chordin* in the dorsal marginal region of the late blastula (Fig. S3), indicating that Fgf2 signalling was indeed active during the late blastula stage. Maternal Fgf2 also possibly triggered *fgf8* activation at this stage, and autonomous *fgf8* expression was reinforced by a positive loop of Fgf signalling, as inferred from the suppression of *fgf8* in Fgf8 morphants (Fig. 6B). Our results correspond with an earlier report that phosphorylated Mapk, an indicator of active Fgf signalling, is located in the dorsal-animal region during the blastula stage (Schohl and Fagotto, 2002). These findings are consistent with the idea that autonomous Fgf signalling via the Mapk/Ets route contributes to initiating neural induction, assuming the onset of *sox2* expression at the blastula stage 9 as the onset of neural induction. Notably, autonomous action of Chordin, Noggin, and Nordal3.1 within a group of cells in the dorsal-animal region is required to initiate neural induction during the late blastula stage (Kuroda et al., 2004). Consistently, activated (phosphorylated) Smad1 expression is maintained at a low level in this period within the dorsal-animal region (Schohl and Fagotto, 2002). Thus, the integration of autonomous positive-Fgf signalling and anti-Bmp signalling in ectoderm cells might be essential for neural induction from the very start.

Besides Fgf2 and Fgf8, Fgf4 is involved in neural induction (Marchal et al., 2009), but *fgf4* is activated around stage 10, with its transcript levels considerably lower than those of Fgf2 and Fgf8 (Suzuki et al., 2017). However, comparison of dose-response profiles of these Fgfs for the activation of neural genes indicated that Fgf4 was more potent than Fgf8 (Fig. S2), revealing the possibility that Fgf4 expressed in the organiser region reinforced neural induction initiated by Fgf2 and Fgf8 and subsequently contributed to neural patterning. It should also be noted that *igf2*, encoding another candidate of neural inducer (Pera et al., 2003), is activated after stage 10 and may contribute to neural induction and patterning together with Fgf4.

### Fgf/Ets signalling is required for neural patterning as an early morphogenic factor

In cultured isolated ectoderm cells, none of the posterior neural genes is activated, but Fgf can induce them to express position-specific neural marker genes along the AP axis in a dose-dependent manner, with lower doses eliciting more anterior genes and higher doses eliciting more posterior genes (Kengaku and Okamoto, 1995). Here, we further showed that Fgf/Ets signalling could dose-dependently activate a different set of position-specific markers, such as *otx2*, *hes7.1*, *foxb1*, and *cdx4* (Fig. 7). Notably, their expression in normal development starts around stage 10 and peaks around stage 12 (Session et al., 2016), coinciding with the establishment of a gradient of the level of phosphorylated Ets within the gastrula ectoderm layer, which increases towards the organiser, located posteriorly (Schohl and Fagotto, 2002). These findings imply that Fgfs could function as early neural patterning morphogens in normal development via the Fgf/Ras/Mapk/Ets pathway in the ectoderm layer. Indeed, the Fgf/Ets route was shown to directly activate *cdx4* (Haremaki et al., 2003), which triggers the expression of several *hox* genes, regulating posterior development during later developmental stages (Isaacs et al., 1998).

Wnt/β-catenin signalling has been reported to regulate the AP neural patterning in *Xenopus* (Kiecker and Niehrs, 2001). However, the gradient pattern of this signalling was not apparent until mid-gastrula stage 11 (Schohl and Fagotto, 2002), suggesting that Wnt/β-catenin signalling may reinforce and further promote the process of neural patterning. Retinoic acid (RA) is another candidate for a neural patterning morphogen. A gradient of RA is established around the stage 12 *Xenopus* neurula, exhibiting a peak at the hindbrain–spinal cord boundary (Pera et al., 2014). Furthermore, RA-responsive enhancers are located at the 3′-region of the *hox* clusters (Langston et al., 1997), which regulate AP patterning in the hindbrain and the spinal cord during a later stage of neural patterning (Strate et al., 2009). Our results, in line with previous research, indicated that Fgfs functioned as morphogens to provide a basis for neural patterning via the Fgf/Ras/Mapk/Ets pathway.

### Source of Fgfs for neural patterning

We attempted to ascertain how the neural patterning Fgfs are derived and contribute to the establishment of the active Ets gradient. *Fgf8* was autonomously activated in cultured ectoderm cells by a positive feedback loop (Fig. 6B). However, this autonomous *fgf8* expression was suppressed by increasing the dose of Fgfs (Fig. S2H), suggesting that negative feedback regulation of *fgf8* expression was also present in ectoderm cells. Our findings are consistent with prevalent observations, including ours, that autonomous neuralization of ectoderm cells is never accompanied by the expression of posterior neural genes (Gruntz and Tacke, 1989; Sato and Sargent, 1989; Hongo et al., 1999; Muñoz-Sanjuán and Brivanlou, 2002). In contrast to the expression of *fgf8* and anterior neural genes, the expression of posterior neural genes required Fgfs in a dose-dependent manner along the AP axis; the higher dose elicited more posterior genes (Fig. 3F and 7E). We previously showed that ectoderm cells, when co-cultured with organiser cells, express position-specific posterior neural genes such as *egr2/krox20*, *hoxc6/XlHbox1*, *hoxc9/XlHbox6*, and *cdx4/Xcad3* in an organiser cell number-dependent manner, which was suppressed by ΔxFgfr-4a overexpression in ectoderm cells (Hongo et al., 1999). These findings imply that, in normal development, organiser cells promote the patterning of the neighbouring neuroectoderm along the AP axis by releasing Fgfs in a paracrine manner. A gradient of active Fgf signalling along the AP axis is established in the neuroectoderm during early (stage 10) to mid (stage 10.5) gastrula stages, with increasing levels towards the organiser region (Schohl and Fagotto, 2002). When we re-examined this aspect, a gradient of P-Mapk localisation along the AP axis in the prospective neural region of gastrula embryos was confirmed (Fig. S4).

### Evolutionary aspects of Fgf/Ets signalling

We wonder how far Fgf/Ras/Mapk/Ets signalling is conserved and plays an essential role in neural development during animal evolution. The requirement of Fgf signals for neural induction is conserved between several vertebrate species, such as zebrafish (Furthauer et al., 2004), *Xenopus* (Hongo et al., 1999; Delaune et al., 2005), and chicken (Streit et al., 2000; Wilson et al., 2000), but the employment of the Fgf/Ets pathway has only been shown for *Xenopus* (this study). Notably, however, two species of ascidians, which share the last common ancestor with vertebrates in chordate evolution, initiate neural induction via the Fgf/Ras/Mapk/Ets pathway, as shown by *otx* expression (Bertrand et al., 2003 in *Ciona intestinalis*; Miya and Nishida, 2003 in *Halocynthia roretzi*). Furthermore, in *Ciona*, the regulatory element driving *otx* expression in the prospective neural cells was identified, which directly responded to Fgf signalling. These findings suggest that the last common ancestor of ascidians and vertebrates in the chordate lineage employs Fgf/Ets signalling for the process of neural induction. It will be interesting to ask whether vertebrate species other than *Xenopus* employ this signalling pathway.

The Fgf/Ras/Mapk/Ets signalling is also involved in neural patterning in *Ciona*, as revealed by *six3* and *otx* expression (Gainous et al., 2015). Remarkably, even some protostomes, such as annelids, arthropods, and onychophorans, express *six3* and *otx* in this order along the AP axis in their anterior neuroectoderm (Steinmetz et al., 2010), as shown in *Xenopus* and *Ciona*, which are classified as deuterostomes. Whether the expression pattern of *six3* and *otx* in protostomes is regulated via Fgf/Ets signalling is yet to be addressed. However, in the planarian, another protostome, Fgfr-related genes were involved in neural development (Cebria et al., 2002). It is tempting to assume that the last common ancestor of bilaterians already acquired the Fgf/Ets signalling pathway for neural development, although the original pathway might have been modified or lost in some of its descendants.

### Concluding remarks

Among a number of candidate signal molecules, Fgf2 and Fgf8 primarily contribute to initiating neural induction. They function in ectoderm cells via the Mapk/Ets route directly at the level of transcriptional regulation of the target genes. This is independent of Bmp signal inhibition via the Mapk/Smad1 route, which was proposed to be the prevailing Fgf signalling route in neural induction. Our findings satisfy the requirement of a different route for neural induction, hypothesized from different lines of evidence. The Fgf signals for neural induction are derived primarily from ectoderm cells themselves in an autonomous manner. This might conflict with the leading idea that the organiser plays the primary role for both neural induction and patterning. However, there is a possibility that the precursors of the organiser cells are present in the dorsal marginal zone of the late blastula apart from regular ectoderm cells, and these precursor cells release Fgfs, contributing to the earliest stage of neural induction in a paracrine mode; the issue is yet to be addressed. Fgfs derived from the organiser at the gastrula stage contribute to neural patterning in a paracrine manner. However, the Fgf members involved were not identified, since loading MOs into the organiser cells blocked their differentiation, disturbing the gastrulation process. Our work provides support for a shared use of Fgf/Ets signalling for neural development during chordate evolution, but further work is needed to ascertain its wider conservation during animal evolution.

## MATERIALS AND METHODS

### Animal care

*Xenopus laevis* embryos were obtained (Mitani and Okamoto, 1989) and staged (Nieuwkoop and Farber, 1967) as previously described. The handling of animals was carried out in accordance with nationally prescribed guidelines and the guidelines for animal experiments at Gakushuin University.

### Plasmid construction

Full or partially deleted coding sequences of *Xenopus*, *ets1* (*xets1*) and *etv1* (*xetv1*), and human *elk1* (*helk1*) were subcloned into pSP64T as described previously (Haremaki et al., 2003); *Δhelk1* RNA blocked Ets transcriptional activity more efficiently than *Δxets1* or *Δxetv1* RNA, albeit being of human origin. *Δhelk1*·*enr* was generated by in-frame C-terminal fusion of the *Drosophila engrailed* repressor region (EnR; Conlon *et al*., 1996) to *Δhelk1*, while *enr*·*Δxetv1* was generated by in-frame N-terminal fusion of EnR to *Δxetv1*. *ΔΔxetv1* is deleted further in the N-terminal region of *Δxetv1* to eliminate its DNA-binding capacity (Papoutsopoulou and Janknecht, 2000), thereby serving as a negative control construct. Structural features of the Ets proteins and their dominant-negative derivatives are shown in Fig. 1A.

### Antisense MOs

Translation-blocking antisense MOs were obtained from Gene Tools LLC (Philomath, OR, USA). These are listed in Supplementary information Table S1.

### Whole-mount *in situ* hybridisation

*In situ* hybridisation experiments were performed following the previously described methods (Harland, 1991; Sive *et al*., 1995). Digoxigenin-labelled antisense RNAs were prepared by the transcription of respective plasmids carrying the target genes, which were linearised. Hybridised digoxigenin-containing RNAs were visualised with anti-digoxigenin antibodies conjugated to alkaline phosphatase and BM purple. For visualising the lineage tracer *gfp* RNA, embryos stained with BM purple were collected in MEMFA (100 mM MOPS (pH 7.4), 2 mM EGTA, 1 mM MgSO_4_, 3.7% formaldehyde) and rinsed twice with PBST (PBS + 0.1 % Tween 20) for 1h. Embryos were then processed for *in situ* hybridisation to *gfp* RNA as described above, except for the use of fluorescein-labelled antisense RNA, anti-fluorescein secondary antibody, and Fast Red. In the experiments described in Fig. 2, stained embryos were bisected as described by Sudou et al. (2012).

### Microinjection of RNAs and Fgf MOs

Capped synthetic RNAs for microinjection were synthesized as described previously (Hongo et al., 1999) and injected into animal blastomeres at the 8-cell, 16-cell, or 32-cell stage. Fgf MOs with or without cognate RNA or wild-type *xetv1* RNA were injected into animal blastomeres at the 8-cell stage. The cognate *fgf2* and *fgf8* RNAs were designed to mismatch with the respective antisense MOs but conserved the same amino acid sequence as the wild-type RNAs. They were synthesized from mutated plasmids that were constructed from wild-type *fgf2* and *fgf8* plasmids via PCR, using 5′ forward primers with mutated cognate sequences (Table S2).

### *Xenopus* ectoderm cell microculture and quantitative RT-PCR

Methods for culturing early gastrula cells were essentially as described previously (Mitani and Okamoto, 1991; Kengaku and Okamoto, 1993; 1995). Animal cap fragments were dissected from injected or uninjected embryos (stage 10), which were dissociated by incubation in Ca^2+^- and Mg^2+^-deficient MBS containing 1% BSA at room temperature. The dispersed cells were then suspended in standard MBS containing 1% BSA, and the desired number of cells were inoculated onto Terasaki plates (150 ectoderm cells/well). After completion of reaggregation by brief centrifugation, cells were incubated in the presence or absence of Fgfs at 22.5°C in humidified air until control embryos reached stage 14 or 25 (Fig. 3A). In some experiments, inhibitors of the Fgf/Ras/Mapk pathway, such as SU5402 (CAS 215543-92-3, Calbiochem) and U0126 (CAS 109511-58-2, Calbiochem), were added with Fgfs. Recombinant Fgfs used were as follows; bovine Fgf2 (bFgf) from Progen Biotechnik GmbH (Heidelberg, Germany), human Fgf4 from PeproTech Inc (Cranbury, NJ, USA), and mouse Fgf8 from R&D Systems Inc (Minneapolis, MN, USA). RT-PCR was performed as described previously (Hongo et al., 1999). PCR products were separated on a 4% polyacrylamide gel, and the radioactivity of each PCR product was estimated using a laser image analyser (Fujix BAS 2000, Fuji Film). When isotopic measurement was not available, the intensity of the band stained in the gel was estimated using ImageJ. PCR products were standardised against a co-amplified internal control, *elongation factor 1α* (*ef1α*; Krieg et al., 1989). The primers used are listed in Table S1.

### Staining of activated Mapk (dpERK)

Whole-mount dpERK staining was performed as previously described (Christian and Slack, 1999).

## ACKNOWLEDGEMENT

We thank Professor Takashi Adachi-Yamada for critical reading of the manuscript.

