## supplemental information for "Fgf/Ets signalling in *Xenopus* ectoderm initiates neural induction and patterning in an autonomous and paracrine manners"

**SUPPLEMENTARY INFORMATION**

**Table S1**

Antisense oligonucleotides and cognate RNAs

| | Sequences |
|---|---|
| FGF2 | |
| peptide | N -MetAlaAlaGlySerIleThrThrLeu•••-C |
| mRNA sense strand | 5'-ATGGCGGCAGGGAGCATCACAACTCTG•••-3' |
| mutated cognate RNA strand | 5'-ATGGCCGCTGGCTCGATAACTACATTG•••-3' |
| antisense oligonucleotide | 5'-GAGTTGTGATGCTCCCTGCCGCCAT-3' |
| 5mis-oligonucleotide | 5'-GAGTTCTGATCCTCGCTGGCGGCAT-3' |
| FGF3 | |
| antisense oligonucleotide | 5'-GCAACAGGATCCAAATTATAACCAT-3' |
| FGF4 | |
| antisense oligonucleotide | 5'-CACCAGGGCCGATGGAACAGTCATC-3' |
| FGF8 | |
| peptide | N -MetAsnTyrIleThrSerIleLeuGly•••-C |
| mRNA sense strand | 5'-ATGAACTACATCACCTCCATCCTGGGC•••-3' |
| mutated cognate RNA strand | 5'-ATGAATTATATAACGAGCATACTCGGC•••-3' |
| antisense oligonucleotide | 5'-CCAGGATGGAGGTGATGTAGTTCAT-3' |
| 5mis-oligonucleotide | 5'-CCAGCATCGAGGTCATGTACTTGAT-3' |
| FGF9 | |
| antisense oligonucleotide | 5'-TGCCAACTTCACCCAGGGGAGCCAT-3' |

**Table S2**

Oligonucleotide primers for RT-PCR

| Markers | Sequences |
|---|---|
| *bf1* | F 5'-TCAACAGCCTAATGCCTGAAGC-3' |
| | R 5'-GCCGTCCACTTTCTTATCGTCG-3' |
| *rax* | F 5'-ACAGCCTTGCAGAGCTTACC-3' |
| | R 5'-CAAGGCTTGCCAATAAACTGG-3' |
| *pax6* | F 5'-TTACCCAGGAACAAATAGAGGCG-3' |
| | R 5'-TGGAACCCGATGTGAATGAGG-3' |
| *en2* | F 5'-GAATAAGCCATCAGGATGAGC-3' |
| | R 5'-GGTCTGTCTGAATATCTGGTGC-3' |
| *hoxc6* | F 5'-TGCAGGCAGAACTCAATGG-3' |
| | R 5'-AAGTGCATTGGCGATCTCG-3' |
| *cdx4* | F 5'-ACCAGCAATAACCACACAGCG-3' |
| | R 5'-TGTAAAATCCCAGTCCCAGATGG-3' |
| *sox2* | F 5'-ACATGAACAGGTCGCCTACC-3' |
| | R 5'-GATAGTGTTGCGACATGTGC-3' |
| *otx2* | F 5'-GGATGGATTTGTTGCACCAGTC-3' |
| | R 5'-CACTCTCCGAGCTCACTTCTC-3' |
| *fgf8* | F 5'-AGTTCTCGGTAGGACACGGTTTC-3' |
| | R 5'-CCCTGCTTGAGGTTCTTCTTCC-3' |
| *hes7.1* | F 5'-CAGCCCTACAAAGACTCTGGTTGAC-3' |
| | R 5'-TCTGTGCCTGCGTGATACAAGC-3' |
| *foxb1* | F 5'-ATCTCCACAGACAGCCACAAGC-3' |
| | R 5'-TGACCACAGTTTGGGAAAGACG-3' |
| *six3* | F 5'-CCTCCAGTGACTCTGAATGTGACG-3' |
| | R 5'-GGTGAGATGAGTGGGATTCCTCTTC-3' |
| *ef1α* (1) | F 5'-GACAACATGCTTGAGCCTAGC-3' |
| | R 5'-CACGACCAACTGGTACAGTACC-3' |
| *ef1α* (2) | F 5'-GGAAAGTCCACAACAACTGG-3' |
| | R 5'-GGAGCATCAATGATAGCGAC-3' |

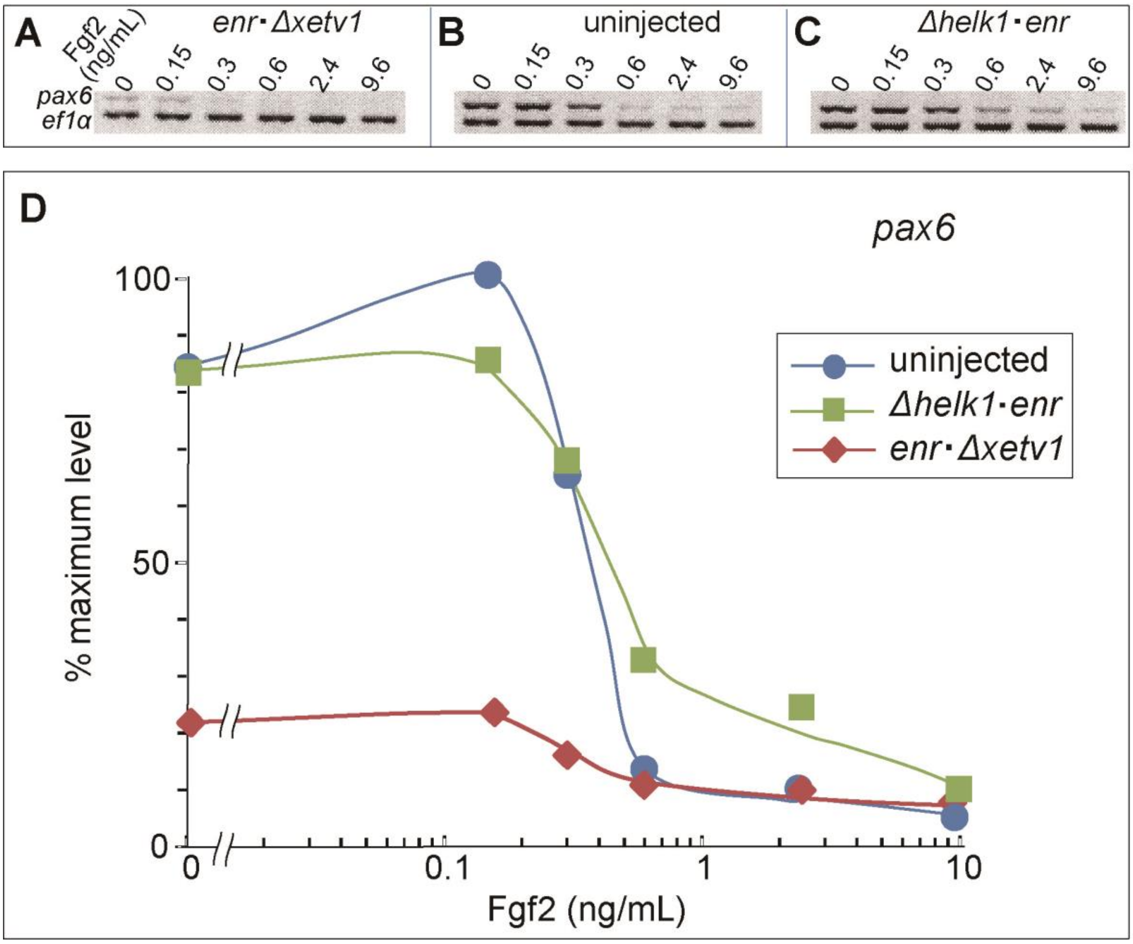

**Fig. S1. Suppression of *pax6* expression in cultured ectoderm cells by blocking Ets transcriptional activity**

(A–C) The experimental design is as described in Fig. 7B–D.

(D) Quantification of the data in (A), (B), and (C).

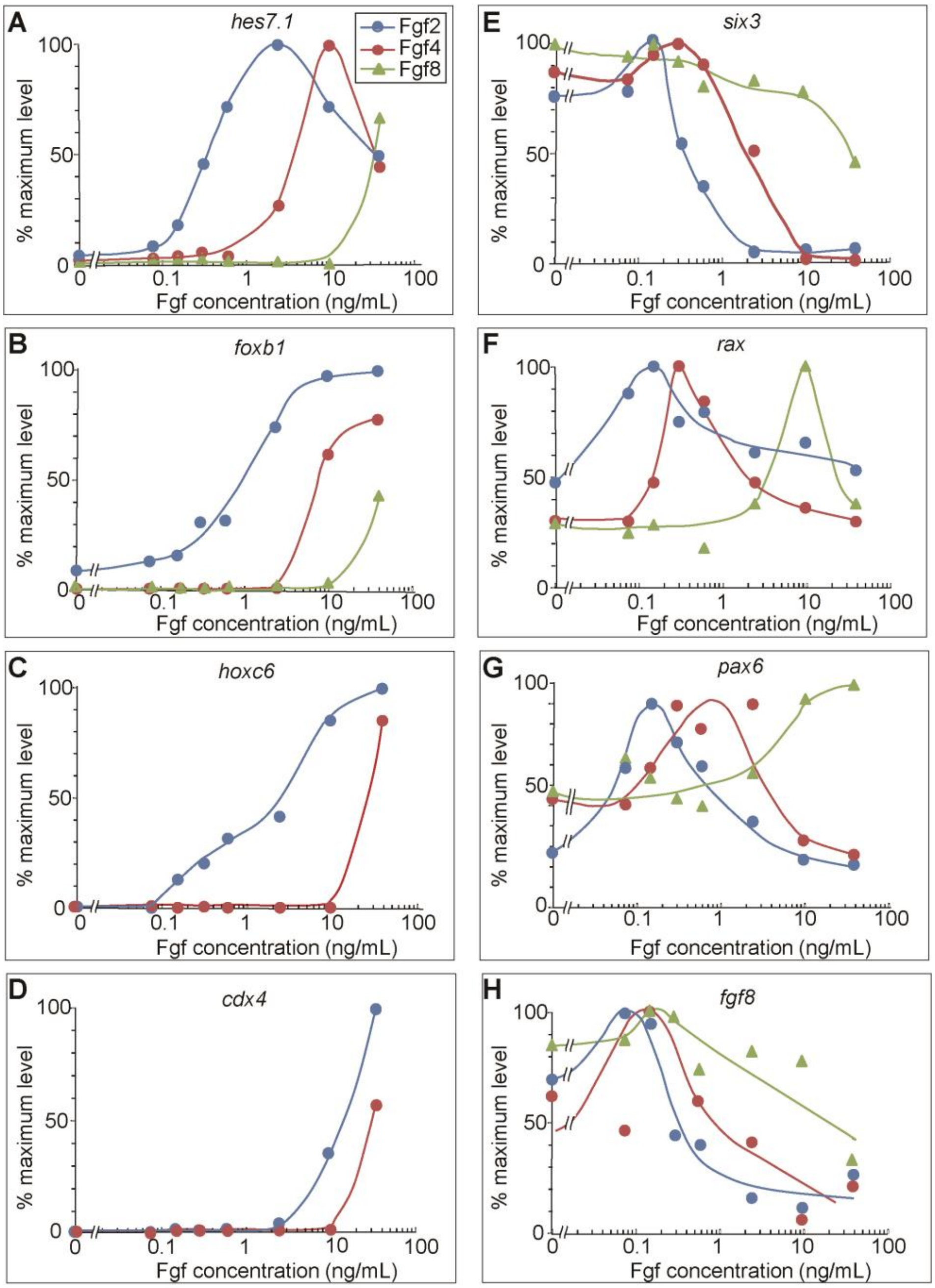

**Fig. S2. Quantitative comparison of Fgf2, Fgf4, and Fgf8 dose-response profiles on the transcript levels of the position-specific neural marker genes and *fgf8* in cultured ectoderm cells.**

The experimental design is as described in Fig. 3, except that the cells were prepared solely from uninjected embryos. The ratio of the band intensities of the genes relative to that of *ef1α* was calculated and plotted against respective Fgf doses. The percentage of the maximum value of the ratio is presented in each profile.

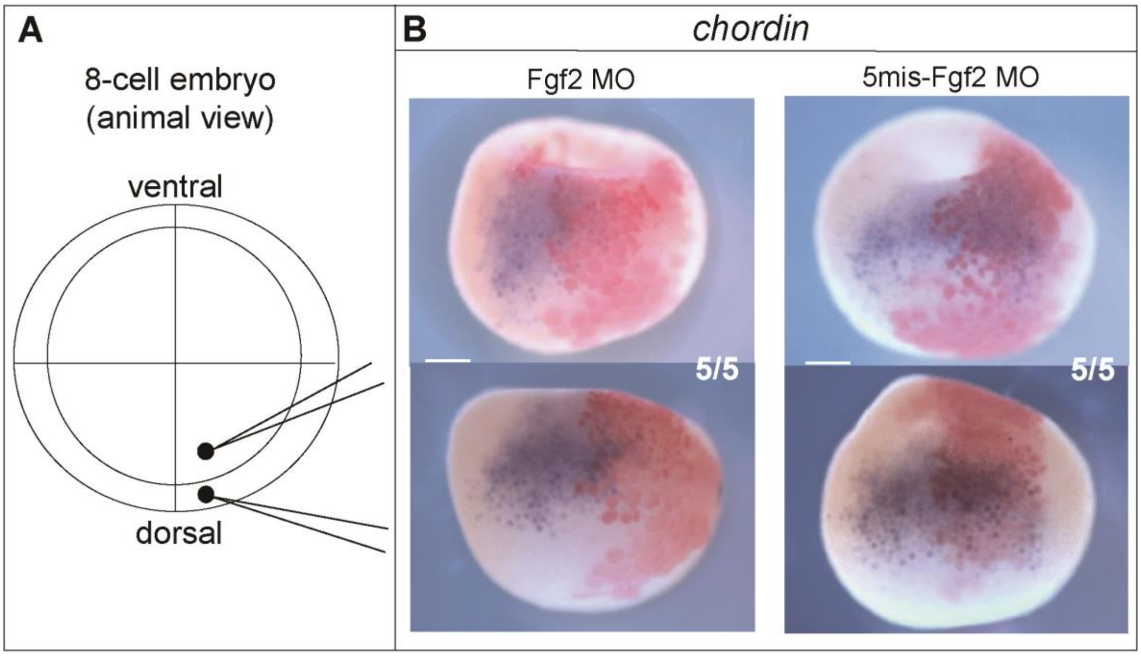

**Fig. S3. Suppression of *chordin* expression in the late blastula via the depletion of Fgf2 in the dorsal marginal zone.**

(A) Injection protocol.

(B) Fgf2 MO was injected into a dorsal-animal blastomere (6.0 ng/blastomere) and a dorsal-vegetal blastomere (12.0 ng/blastomere) at the 8-cell stage. Injected embryos were cultured until stage 9.5 and processed for *in situ* hybridisation. Scale bar = 0.25 mm.

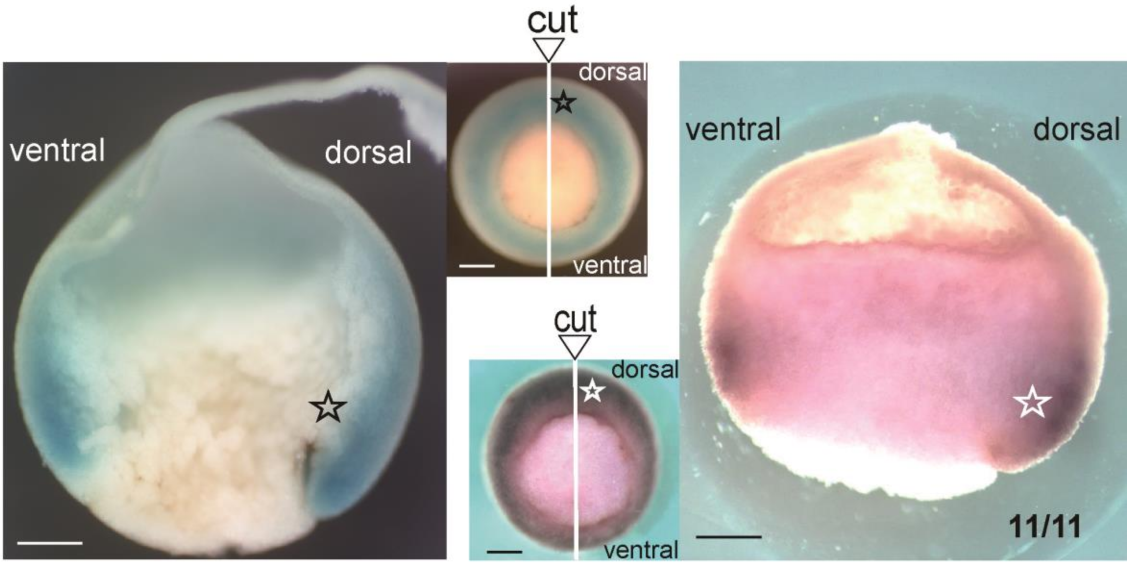

**Fig. S4. The gradient of the activated (phosphorylated) Ets transcription factor within the gastrula ectoderm layer.** Whole-mount phosphorylated Ets (dpERK) staining was performed as described previously (Christian and Slack, 1999). Vegetal (middle panels) and cut-surface views of the two typical results are shown.

☆ indicates the dorsal right side of each embryo. Scale bar = 0.25 mm.
